# Embryonic Exposure to Valproic Acid impairs Social Predispositions for Dynamic Cues of Animate Motion in Newly-Hatched Chicks

**DOI:** 10.1101/412635

**Authors:** Elena Lorenzi, Alessandra Pross, Orsola Rosa-Salva, Elisabetta Versace, Paola Sgadò, Giorgio Vallortigara

## Abstract

Predispositions to preferentially orient towards cues associated with social partners, such as face-like stimuli or biological and animate motion, appear to guide social behavior from the onset of life. These predispositions have been documented in several vertebrate species including human neonates, young monkeys and newly-hatched domestic chicks. Human newborns at high familiar risk of Autism Spectrum Disorder (ASD) show a remarkable deficit in their attention toward these predisposed stimuli, either static and dynamic, compared to newborns at low risk. A previous study showed that prenatal exposure to valproic acid (VPA) (that in humans increases the risk of developing ASD) impairs the chicks’ predisposition to approach naturalistic social stimuli that convey static configurations of features (a stuffed hen), without impairing general cognitive and learning abilities. Here we investigated the effect of VPA exposure on another class of social predispositions, *i.e.* the spontaneous preference to approach self-propelled objects, namely objects that display autonomous changes in speed. We observed that the preference for stimuli displaying autonomous changes in speed was impaired in VPA-compared to vehicle-injected chicks. Our results indicate an effect of VPA on the development of predisposed orienting mechanisms towards dynamic stimuli, that could be used to investigate the molecular and neurobiological mechanisms underlying ASD early symptoms.

## Introduction

Neonates of some vertebrate species display unlearned, rudimental knowledge that orients their early social responses towards objects with the same features present in social partners and caregivers. Comparative research on human infants and newly-hatched domestic chicks (*Gallus gallus*) found striking similarities in the static and dynamic visual cues that attract their attention soon after birth (Di Giorgio et al., 2017). Among static cues, a predisposition to approach a stuffed hen over a scrambled version of the same stimulus has been consistently observed in visually-naïve chicks (Bolhuis and Horn, 1997; Johnson and Horn, 1988; Mayer et al., 2016; Versace et al., 2017). This preferential approach is driven by the configuration of features present in the intact head and neck (Johnson and Horn, 1988). Similarly, schematic face-like configurations (three dark blobs triangularly arranged within an oval outline, two in the eyes region and one in the nose-mouth region) attract human neonates and visually-naïve chicks (Farroni et al., 2005; Rosa Salva et al., 2010; 2011; Turati et al., 2002). Among dynamic cues, point-light displays depicting biological motion are preferred by neonates of both species to the same configuration of dots rigidly rotating or moving randomly (Simion et al., 2008; Vallortigara and Regolin, 2006; Vallortigara et al., 2005). Moreover, simple shapes that move in a self-propelled fashion are preferred over objects that do not exhibit this property (Mascalzoni et al., 2010; Rosa-Salva et al., 2016). Chicks seem to have a spontaneous preference for objects autonomously starting to move over objects set on motion by a collision (Mascalzoni et al., 2010) and for objects autonomously changing their speed over constant moving ones (Rosa-Salva et al., 2016). Similarly, human neonates exhibit a looking preference for self-propelled objects autonomously starting from rest (Di Giorgio et al., 2016b). The adaptive function of these predispositions has been hypothesized to be in directing attention toward highly important animate stimuli, enabling future learning through experience (Di Giorgio et al., 2016a; 2017; Versace et al., 2018) and enhancing social interactions (Powell et al., 2018). In chicks, predispositions are likely to orient the young animal toward the mother hen (or other brood mates), directing subsequent filial imprinting responses towards animate stimuli, and thus maintaining the brood cohesion and enabling social learning from conspecifics (Miura and Matsushima, 2016; Rosa Salva et al., 2015). For human neonates, selective attention towards animate beings, such as the caregiver, is even more crucial, since altricial human infants are completely dependent on maternal care (Di Giorgio et al., 2017).

Alterations in social predispositions appear to be linked to Autistic Spectrum Disorders (ASD). ASD comprise a complex group of neurodevelopmental disabilities characterized by a wide range of impairments, including important deficits in the domain of social cognition (Sacrey et al., 2015). Impairments in face discrimination and recognition have been widely observed in ASD individuals (Dawson et al., 2005). Similarly, altered processing of stimuli depicting biological motion has been reported (Freitag et al., 2008): two-year-olds with ASD do not orient towards biological motion point-light displays whereas they are attracted to non-social physical contingencies (Klin et al., 2009). In line with the hypothesis of an impairment in the unlearned social predispositions, young children (48 months old) with ASD show difficulties compared to control children in spontaneously categorizing self-propelled motion as animate (Rutherford et al., 2006). Interestingly, after a training period, these children are able to learn the categorization task, an outcome that emphasizes the difference between unlearned knowledge and experience. Early differences have been also observed between neonates at high- and low-risk for ASD in their visual attention toward faces and biological motion (Di Giorgio et al., 2016a). Neonates at high familiar risk of ASD show significant differences compared to low-risk neonates in the preference for a face-like stimulus and for biological motion, suggesting an impairment or a delay in the development of the spontaneous mechanisms detecting animate beings. Observing the same impairment for both static and dynamic stimuli in a different species would argue in favor of a common developmental origin of these predispositions.

In humans, prenatal exposure to valproic acid (VPA) has been shown to increase the risk of developing ASD (Christensen et al., 2013) and embryonic exposure to it has been widely used to model the ASD syndrome in rodents (Choi et al., 2016; Morris et al., 2010; Nagode et al., 2017; Schneider and Przewłocki, 2005). Embryonic exposure to VPA has been shown to induce impairments in chicks’ aggregative behavior (Nishigori et al., 2013). Moreover, chicks exposed to VPA during embryonic development are impaired in their early predisposition for static stimuli (Sgadò et al., 2018). Dark-reared VPA-injected chicks show no spontaneous preference to approach a stuffed hen over a scrambled version of the same stimulus (Sgadò et al., 2018). Interestingly, these studies with chicks found that filial imprinting, an experience-dependent learning mechanism, was unaffected by VPA-injection, supporting the hypothesis of a selective impairment in unlearned predispositions (Nishigori et al., 2013; Sgadò et al., 2018).

To further study the effect of VPA on early predispositions, and whether the impairment of predispositions for static configurations is accompanied by impairment in predispositions for dynamic stimuli, we compared the spontaneous preference to stimuli conveying self-propulsion cues in VPA- and vehicle-injected chicks.

## Materials and Methods

### Ethical statement

All experiments comply with the current Italian and European Community laws for the ethical treatment of animals. The experimental procedures were approved by the Ethical Committee of the University of Trento and licensed by the Italian Health Ministry (permit number 986/2016-PR).

### Embryonic injections

Fertilized eggs of domestic chicks (*Gallus gallus*), of the Ross 308 (Aviagen) strain, were obtained from a local commercial hatchery (Agricola Berica, Montegalda (VI), Italy) and incubated at 37.7 °C and 60% of relative humidity in the darkness. The first day of incubation was considered embryonic day 0 (E0). At E14, fertilized eggs were selected by candling before injection. Embryo injection was performed according to previous reports (Nishigori et al., 2013; Sgadò et al., 2018). Briefly, a small hole was made on the eggshell above the air sac, and 35 μmoles of VPA (Sodium Valproate, Sigma Aldrich) were administered to each fertilized egg, in a volume of 200 μl, by dropping the solution onto the chorioallantoic membrane. Age-matched control eggs were injected using the same procedure with 200 μL of vehicle (double distilled injectable water). After sealing the hole with paper tape, eggs were placed back in the incubator (FIEM srl, Italy). During incubation and hatching, eggs and chicks were maintained in complete darkness, preventing any visual experience prior to the test. The day of hatching each chick was tested only once.

### Apparatus, stimuli and test

We used the same procedure previously described (Rosa-Salva et al., 2016) to assess chicks’ predispositions for speed-change. Briefly, carefully avoiding any other visual experience, the day of hatching chicks were individually placed in the center of the test apparatus, a corridor (85×30×30 cm), open at the two ends where two video screens were displaying the experimental stimuli. The corridor was divided in three sectors: a central sector (45 cm long) delimited by two steps, that the animals had to climb to enter the two choice sectors (each 20 cm long) immediately adjacent to the two screens. Stimuli were two video-animations representing the movement of a simple red circle. In one video the object was moving at constant speed (speed-constant) and in the other one it was changing its speed (accelerating and decelerating; speed-change). A spontaneous choice test of six minutes was performed for the two stimuli. Chicks’ preference for the speed-change stimulus was measured by the ratio of time (in seconds) spent in the choice sector near the speed-change stimulus divided by the cumulative time spent in either of the choice sectors. Chicks remaining in the central sector were not included in the analyses. Values of this ratio could range from 0 (full choice for the speed-constant), to 1 (full choice for the speed-change), whereas 0.5 represented no preference. For more detailed information on the procedure, see Rosa-Salva et al., (2016). Chicks’ level of motility was measured by evaluating the latency (in seconds) to first approach irrespective of the stimulus approached.

### Data analysis

Effects of Treatment (VPA- and vehicle-injection) and Sex (male, female) on the preference for the speed-change stimulus were assessed by a multifactorial analysis of variance (ANOVA) on the dependent variable ratio. One-sample two-tailed *t*-tests were run to test significant departures from chance level (0.5) of the ratio, separately for the two groups. The number of chicks that first approached the speed-change or the speed-constant stimulus in the two treatment groups was compared using the chi-square test of independence. Effects of Treatment and Sex on latency to first approach were assessed by an ANOVA on the latency to first approach one of the stimuli. All statistical analyses were performed with IBM SPSS Statistic for Windows (*RRID:SCR_002865*). Alpha was set to 0.05 for all the tests.

## Results

For the present study 51 VPA-injected (males=27) and 52 vehicle-injected (males=26) chicks were tested. Results of the ANOVA on the preference for the speed-change stimulus showed a significant effect of Treatment (F_(1,99)_=4.296, p=0.041; Fig. 1A), and no significant effect of Sex (F_(1,99)_=0.0001, p=0.992) nor any significant interaction (Treatment × Sex: F_(1,99)_=0.151, p=0.698). In the control group (vehicle-injected), the preference for approaching the speed-change stimulus was similar to what previously observed (Rosa-Salva et al., 2016) with a preference score significantly higher than chance level (t_(51)_=2.365, p=0.011; *M*=0.673, *SEM*=0.066, Fig. 1A). On the contrary, VPA exposure significantly reduced the preference for the speed-change stimulus: the preference score for approaching the speed change stimulus did not differ from chance level (t_(50)_=-0.406, p=0.686; *M*=0.472, *SEM*=0.696, Fig. 1A). A significant difference between the two groups was found also in the number of chicks that first approached the speed-change stimulus (X^2^=4.314, p=0.047). While in the vehicle-injected group a significantly higher number of chicks first approached the speed-change stimulus (X^2^=6.231, p=0.018; speed-change N=35, speed-constant N=17), in the VPA-treated group no significant difference was found in the number of chicks that approached the two stimuli (X^2^=0.176, p=0.78; speed-change N=24, speed-constant N=27).

**Figure 1.**
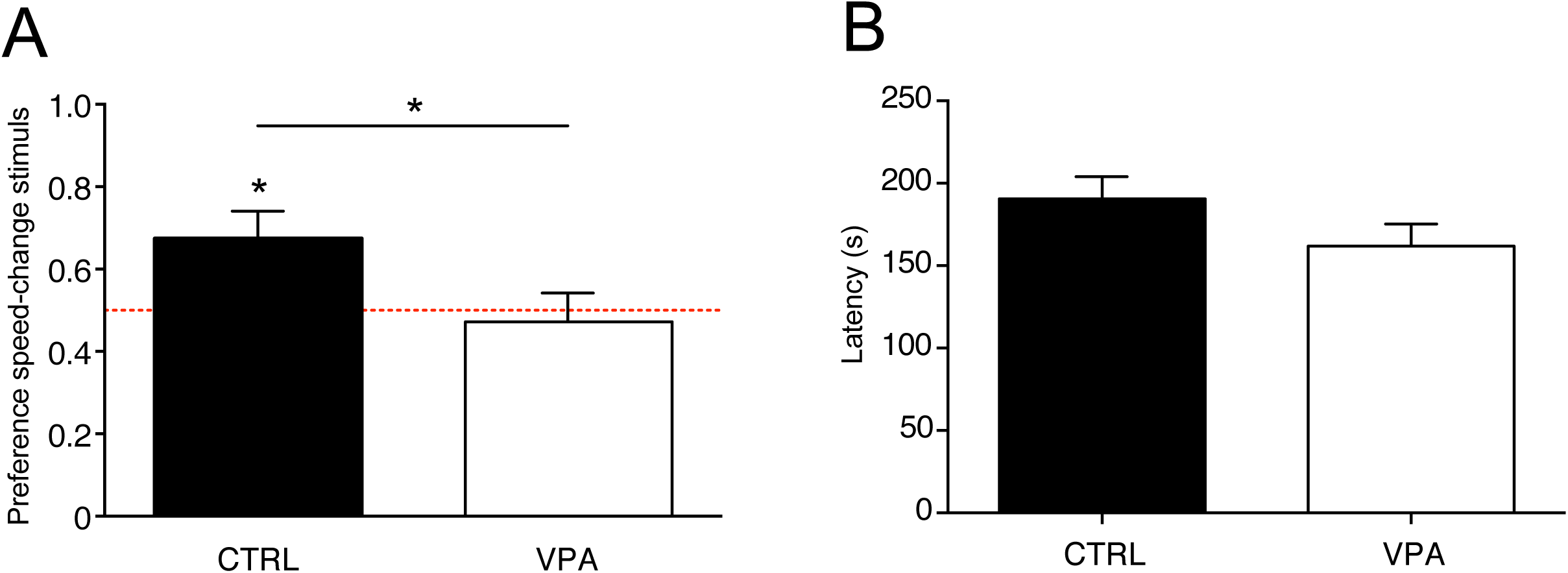
(**A**) Social preference responses for the speed-change stimulus shown as the ratio of time (in seconds) spent in the choice sector near the speed-change stimulus divided by the cumulative time spent in either of sectors (see Methods for details). Analysis of variance of social preference scores using Treatment and Sex as between-subject factors, revealed a significant main effect of Treatment (line with asterisks), with no other main effects or interactions among the other factors analysed. Preference scores were significantly different from chance level for the control group, but not for VPA-treated chicks. Asterisks on top of bars indicate significant departures from chance level, marked by the red line at 0.5. (**B**) Latency to first approach assessed as a measure of motor activity. Analysis of variance on number of rotations using Treatment and Sex as between-subject factors, showing no significant main effects of Treatment, Sex or interaction Treatment × Sex. Data represent mean ± SEM, *p < 0.05.

To evaluate motility, we measured the latency to the first approach, independent of the stimulus, and found no significant effects of Treatment (F_(1,99)_=2.672, p=0.105; Fig. 1B), Sex (F_(1,99)_=1.124, p=0.292), nor any interaction (F_(1,99)_=0.000, p=0.99).

## Discussion

Among the early predispositions described in the human developmental literature, some appear to be impaired in ASD. Human faces and face-like configuration stimuli, a class of static social stimuli, elicit remarkable social attention and orientation responses in typical developing neonates (Simion and Di Giorgio, 2015), while impairments in face and eye-gaze direction processing have been extensively described in infants at risk of ASD (Di Giorgio et al., 2016a; see for a review Webb et al., 2017). Among dynamic stimuli, biological motion and animacy appear to elicit predisposed social-orienting responses in typically developing children that seem to be compromised in infants at high risk of ASD (Di Giorgio et al., 2016b; Klin et al., 2009; Simion et al., 2008). Several accounts suggest that abnormalities in the early social-orienting system may lead to deficits in social stimuli processing, limiting attention to salient social stimuli and decreasing their reward value, resulting in the atypical development of social behavior associated with ASD.

Our data support this hypothesis in an animal model, showing a suppression of predisposed preferences to dynamic as well as static cues to social stimuli in domestic chicks exposed to VPA, a drug used to induce ASD-like behavioral deficits in vertebrate species (Ranger and Ellenbroek, 2016). The observation of a similar impairment in predispositions in different species for both static and dynamic cues suggests a common developmental origin of these social-orienting predispositions. It appears that prenatal exposure to VPA in chicks could be a viable model to investigate the molecular and neurobiological mechanisms underlying those ASD early symptoms that are associated with predisposed orienting mechanisms towards social stimuli.

## Author Contributions

P.S., E.V., O.R.-S. and G.V. conceived and designed the experiments; E.L. and A.P. conducted the experiments; P.S., E.V. and O.R.-S. developed the behavioural paradigms; E.L., A.P., P.S., O.R.-S. and E.V. analyzed the data; E.L. and P.S. drafted the manuscript; E.L., A. P., O.R.-S., E.V., P.S. and G.V. wrote the manuscript. All the authors gave final approval for publication.

## Acknowledgments

We thank Dr Tommaso Pecchia for help with the experimental apparatus, Dr Uwe Mayer for help with the development of apparatus and procedure and Dr Ciro Petrone for animal facility management.

